# Compensatory Relationship between Low Complexity Regions and Gene Paralogy in the Evolution of Prokaryotes

**DOI:** 10.1101/2022.09.23.509281

**Authors:** Erez Persi, Yuri I. Wolf, Svetlana Karamycheva, Kira S Makarova, Eugene V. Koonin

## Abstract

Evolution of genomes in all life forms involves two distinct, dynamic types of genomic changes: gene duplication (and loss) that shape families of paralogous genes and extension (and contraction) of low complexity regions (LCR), which occurs through dynamics of short repeats in protein-coding genes. Although the roles of each of these types of events in genome evolution have been studied, their co-evolutionary dynamics is not thoroughly understood. Here, by analyzing a wide range of genomes from diverse bacteria and archaea, we show that LCR and paralogy represent two distinct routes of evolution that are inversely correlated. Emergence of LCR is a prominent evolutionary mechanism in fast evolving, young protein families, whereas paralogy dominates the comparatively slow evolution of old protein families. Analysis of multiple prokaryotic genomes shows that the formation of LCR is likely a widespread, transient evolutionary mechanism that temporally and locally affects also ancestral functions, but apparently, fades away with time, under mutational and selective pressures, yielding to gene paralogy. We propose that compensatory relationships between short-term and longer-term evolutionary mechanisms are universal in the evolution of life.

**Significance:** Evolution of genomes in all organisms involves a variety of changes occurring on different spatial and temporal scales, from point mutations to whole genome duplication. Here we demonstrate that during the evolution of bacterial and archaeal genomes, there is a universal inverse relationship between the formation of low complexity regions in protein sequences through proliferation of short repeats and gene duplication. The former process apparently is a route of short-term adaptation whereas the latter one dominates evolution on longer temporal scales. We propose that compensatory relationships between evolutionary mechanisms acting at different spatial and temporal scales are a general feature of the process of evolution.

## Introduction

Beyond point mutations, genomes evolve through diverse, dynamic events, in particular, gene gain via duplication and horizontal gene transfer (HGT), and loss, yielding families of paralogs^1,2^. The emergence of gene paralogy is accompanied by relaxed purifying selection, and in some cases, positive selection, such that either new functions arise (neofunctionalization) or existing functions adapt to new conditions and become more specific (subfunctionalization)^3,4,5^. Another prominent phenomenon in genome evolution is the expansion and contraction of low complexity regions (LCR), sections of genes that consist of short repetitive elements, primarily through replication slippage^6^. Typically, LCR manifest as tracks of irregular recurrence of several (or even a single) amino acids in proteins, or of a few (in most case, 1-5) base pairs in nucleotide sequences, such as microsatellites. Although initially perceived as ‘junk’, there is increasing evidence that many LCRs are evolutionarily conserved and functionally important and, with clinical and biotechnological implications^7,8,9^. LCRs have been often linked to intrinsically disordered proteins, which are involved in a variety of functions, notably signaling and regulation^10,11,12^. In particular, LCRs play key roles in regulating the solubility and folding state of proteins^13^. Furthermore, LCR form diverse secondary structures including distinct types of helices and sheets, which are involved in the formation of important protein structures such as collagen, keratin, and cell wall proteins, and are often associated with fast evolution and functional diversification of proteins^14-18^. For example, LCRs are responsible for the formation and mechanical properties of silk proteins^19,20^, provide substrate for the diversification and immune adaptability of antibodies^21,22^, and confer adhesive and self-assembly properties to various proteins^23^. LCRs are highly dynamic on short time scales and can be induced by stress and environmental factors, such as temperature and pH, to regulate protein phase transition. In this capacity, LCRs play important roles in a variety of biological processes, such as amyloidogenesis^24^ and liquid-liquid phase separation, which drives the formation of membrane-less organelles (e.g., nucleolus and stress granules) that regulate numerous cellular functions in response to stress^25-28^ including viral-host interaction^29^. Notably, LCRs are directly involved in many pathologies, in particular, in a range of inherited neurological disorders^30-35^ and in the somatic evolution of numerous cancers^36-39^.

Despite the recognition of the importance of these two routes of evolution, the formation of paralogous gene families and LCRs, the evolutionary interplay between the two (if any), to our knowledge, has not been investigated. It remains unknown whether these distinct evolutionary phenomena occur independently in a stochastic manner, or exhibit some spatio-temporal dependency that could be functionally and evolutionarily important. Insights from studies on cancer evolution suggest that proliferation of short repeats and gene amplification occur in different phases of evolution, with compensatory effects on tumor fitness, with contrasting clinical manifestations^40^. However, it remains unknown whether such a compensatory relationship also plays a role in species evolution. Here we analyze paralogous gene families and LCRs in multiple, diverse genomes of bacteria and archaea, and show that compensation between these two types of genome evolution events is indeed observed universally across prokaryotes.

## Results

### Low complexity regions and gene paralogy in the evolution of groups of prokaryotic species

To study the relationship between the extents of LCR and gene paralogy, we analyzed a set of 8 groups of prokaryotic species, 4 archaeal and 4 bacterial (**Methods**). For each group, we determined the orthologous relationships of proteins across genomes and estimated the levels of LCR and paralogy in each local cluster of orthologous genes (COG) (**Methods**). Several methods have been developed over the years to identify LCRs, devoted primarily to the analysis of protein sequences. The most common include quantifying the irregularity of single amino acids based on Shannon’s entropy, known as SEG algorithm^41^, comparing query sequences against degenerate homopeptides datasets, such as CAST algorithm^42^, graph-based method, such as SIMPLE algorithm^43^, and approaches based on the statistics of motif recurrences^44^. Although all these methods provide comparable results^45,46^, they can differ substantially in many cases, and a consensus is still lacking^18^. To this end, here, LCRs were identified by employing and extending the compositional order approach, which detects irregular recurrences of short *k*-mers^16,39,47^ (see **Methods** for the description and link to the pipeline).

A large fraction of the COGs (30-50%, across the group of species) exhibit neither gene paralogy (PAR=1) nor LCR (LCR=0). This point represents a large ‘attractor’ of functions, largely, house-keeping, essential ones, that evolve without resorting to LCR or paralogy, and obviously, contains no information about the evolution and co-evolution of LCR and paralogy. Excluding this point at the origin, we observed a universal, strong and significant negative correlation between LCR and paralogy (**Figure 1A and Fig. S1**). We analyzed the strength of this correlation in the different functional categories of COGs and found the inverse relationship in most of these. The effect was most pronounced in categories **D** (cell division), **K** (transcription), **L** (replication and repair), **V** (defense and offense systems), **X** (mobilome) as well as among COGs with only generally predicted (**R**) or unknown (**S**) function, the functional categories that have been previously characterized as evolutionarily dynamic, with high rates of gene gain and loss^48^ (**Figure 1B, and Fig. S2**). These observations indicate that the inverse relationship between LCR and paralogy, manifested as a ‘moon’ shape scatter of points (**Figure 1A**,**B**), is a general feature of COGs, universal across species and functions. This pattern suggests that, generally, LCR and paralogy represent distinct, compensatory routes of evolution: that is, either mechanism can promote the evolution and adaptability of biological functions, such that, when one is operational, there is less need for the other.

**Figure 1.**
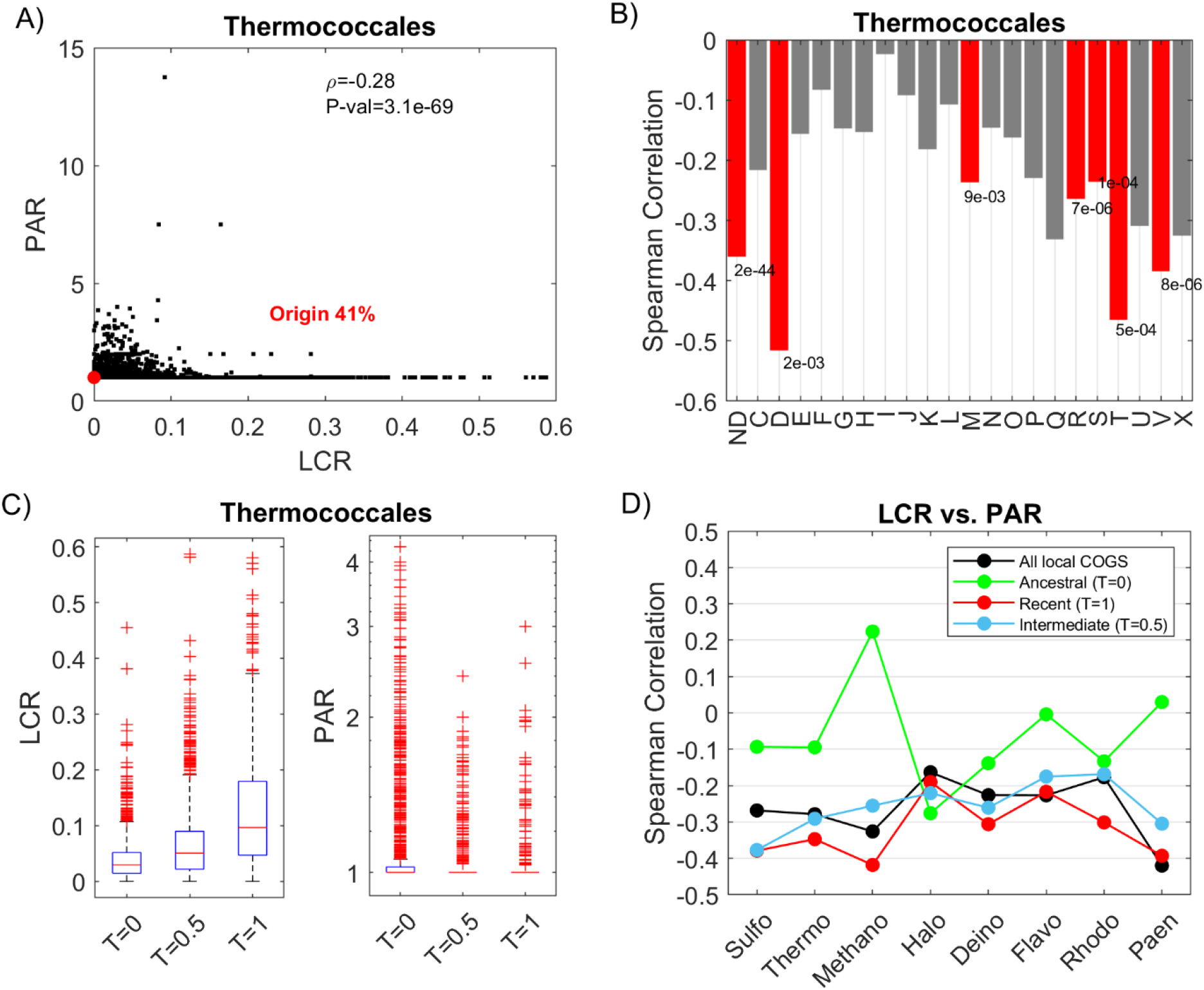
The relationship between low complexity regions and gene paralogy among local COGs within 8 groups of bacteria and archaea. (**A**) Example of the case of the Thermococcales group (n=42), each point is a local COG (n=6450). The COGs at the origin (i.e., LCR=0 and PAR=1) comprise 41% of all COGs (red circle). The Spearman correlation between LCR and PAR, excluding the origin, is −0.28 (P-value=3.1e-69). This inverse relationship between LCR and PAR is apparent across the 8 groups of species (**Fig. S1**) (**B**) Spearman correlation across different functional categories of COGs in Thermococcales. Cases with P-value < 0.01 are colored (red bars). The dominance of this negative correlation across functions is apparent across the 8 groups of species (**Fig. S2**) (**C**) The extent of LCR for locally ancestral COGs (T=0) is significantly lower than that for intermediate (T=0.5) and recently gained COGs (T=1), based on the species tree of Thermococcales. In contrast, PAR tends to be somewhat greater for T=0 than for T>0. These trends are exhibited, in different degrees, across the majority of the 8 group of species (**Fig. S3 and S4**). (**D**) Spearman correlation between LCR and PAR for COGs of different relative age in each of the 8 groups of species. The correlation coefficients are shown for all COGs (black), ancestral COGs (T=0, green), recently evolved COGs (T=1, red), and intermediate age COGs (T=0.5, blue). Abbreviations of functional categories (bold type shows the categories with the most pronounced anticorrelation between LCR and PAR across at least 4 groups of species): ND – not determined, C - energy production and conversion **D - cell division**, E - amino acid metabolism and transport, F - nucleotide metabolism and transport, G - carbohydrate metabolism and transport, H - coenzyme metabolism, I - lipid metabolism, J – translation, **K – transcription, L - replication and repair**, M - membrane and cell wall structure and biogenesis, N - secretion and motility, O - post-translational modification and chaperones, P - inorganic ion transport and metabolism, Q - Secondary metabolism, **R - general functional prediction only, S - function unknown**, T - signal transduction, U - intracellular trafficking and secretion, **V - defense [and offense] systems, X - mobilome**

To elucidate and compare the evolutionary dynamics of LCR and gene paralogy, we constructed species trees from 16S rRNA for the 8 groups of prokaryotes and assigned the relative age to each COG: Ancestral (T=0), if the COG exists in all species, New (T=1), if the COG appears only in one leaf (that is, a singleton or a family of paralogs represented in one member of the group only), or Intermediate (e.g. T=0.5) if the COG is represented in a subset of the species. Generally, LCR is low in ancestral COGs but is high in new COGs, whereas the values for COGs of intermediate age fall in between (**Figure 1C and Fig. S3**). This observation suggests that LCR is a fast, transient mode of adaptation in recently evolved proteins, which tends to subsequently fade away with time and selective pressures. In contrast, paralogy is generally higher in ancestral than in new COGs (**Figure 1C and Fig. S4**), indicating that gene gain by duplication and HGT is a dominant route of evolution in more slowly evolving, ancestral COGs. Given the high paralogy and low or non-existent LCR in ancestral COGs, it could be expected that there should be only weak or no correlation between LCR and paralogy in this case, whereas in new COGs, where LCR is high, one should expect significant negative correlation. Testing this prediction across the 8 groups of species, we indeed found values around zero for ancestral COGs, but strong negative correlation for new COGs, with values close to the reference (all COGs) for COGs of intermediate age (**Figure 1D**).

### Low complexity regions and gene paralogy in individual prokaryotic genomes

We then explored the patterns of LCR and paralogy in single genomes. We hypothesized that large gene families (PAR>1) would typically encompass less LCR, whereas many small families (especially single copy proteins) would resort to LCR as a route of evolution. We tested this conjecture on the 8 groups of species, by analyzing each single genome and reassessing the extent of LCR and paralogy for the COG members and individual proteins that genome encompassed. In this case, PAR is simply the number of COG members (paralogs) in a genome, and the extent of LCR is calculated for all respective proteins. We found that the LCR-PAR anticorrelation in single genomes was substantially stronger than in the previous case, when COGs (and their LCR and PAR values) were analyzed across entire groups of organisms (**Figure 2A-2B** and **Fig. S5**). Surprisingly, in single genomes, we could not separate the LCR-PAR correlation between the ages of COGs as both ancestral and new COGs exhibited strong negative correlation (**Figure 2C** and **Fig. S6**). Examining the potential causes of the unexpected strong correlation in the ancestral COGs, we found that LCR (but not PAR) was much higher in single genomes than when calculated for all organisms in the respective group of species, thereby leading to the strong negative correlation in all COGs, including locally ancestral ones (**Fig. S7**). This observation is the most compelling evidence that LCR is a fast and transient evolutionary mechanism to which ancestral functions also resort on a short time scale, whereas on longer spans of evolution, as time progresses, LCR often fade away under the pressure of purifying selection. Those LCRs that are fixed in the population and survive the pressure of purifying selection over long time spans, in all likelihood, perform a variety of functions.

**Figure 2.**
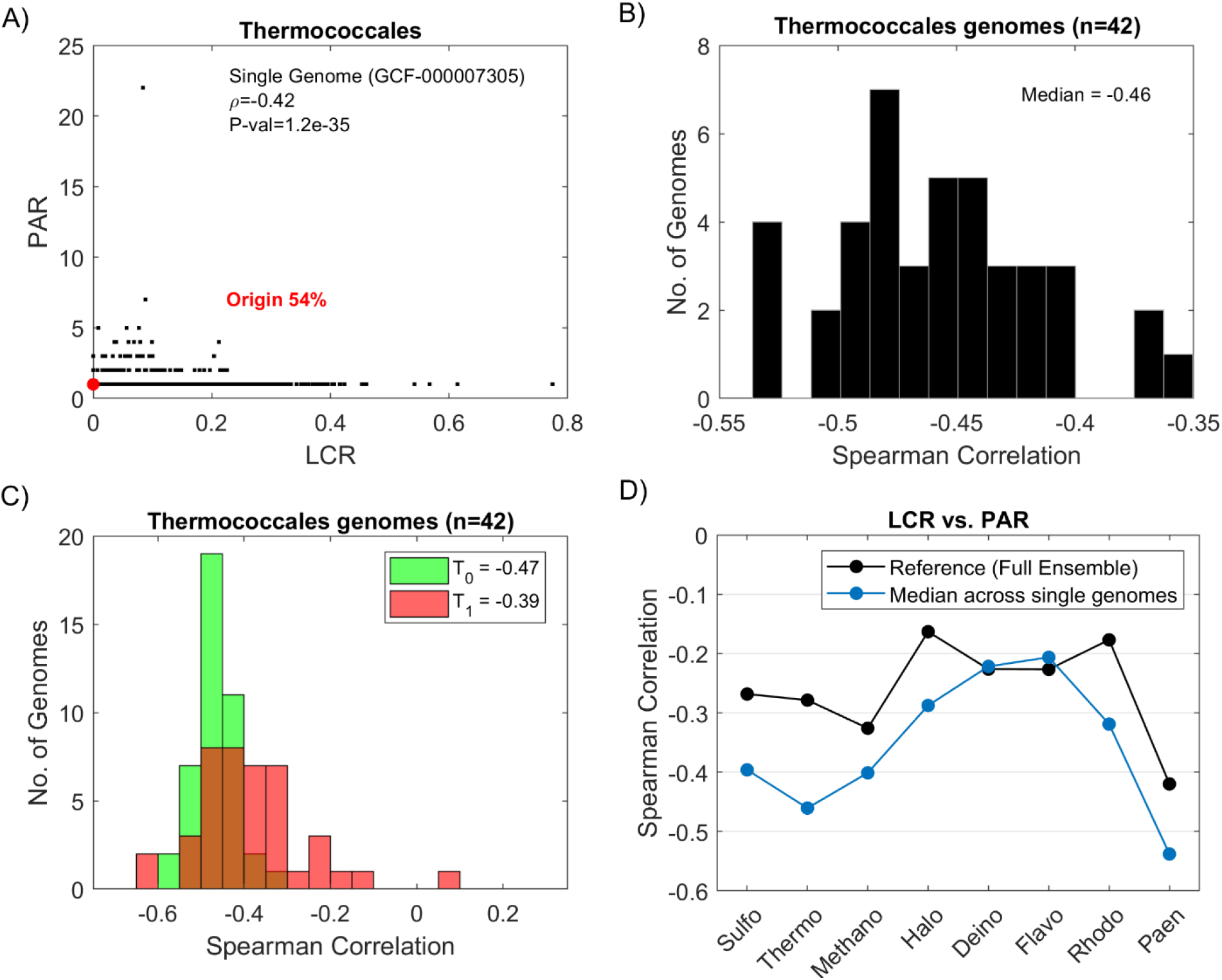
Relationship between low complexity regions and gene paralogy for local COGs in single genomes, across 8 groups of species. (**A**) Example of a single genome of Thermococcales (GCF_000007305.1 - *Pyrococcus furiosus* DSM 3638); each point is a local COG. The points at the origin (i.e., LCR=0 and PAR=1) correspond to 54% of the COGs. The Spearman correlation between LCR and PAR excluding the origin is −0.42 (P-value = 1.2e-35). (**B**) This inverse relationship is observed in all single genomes (n=42) that comprise this group of species. This negative correlation is found in all 8 groups of species (**Fig. S5**). (**C**) The negative correlation is strong for both locally ancestral COG (T=0) and recently gained COGs (T=1). This also occurs in the 8 groups of species (**Fig. S6**). This is a consequence of the much larger LCR for genes associated with a COG in a single genome, compared with the LCR level of this COG for all genes associated with it across all genomes in the given group of species (**Fig. S7**). (**D**) The medians of the distribution of the single-genome Spearman correlation coefficients across genomes (blue), in each of the 8 groups of species, relative to the estimates from **Fig. 1** taken as reference points (black).

## Discussion

In this work, we observed a previously unnoticed, prominent anticorrelation between the extent of LCR and paralogy in prokaryotic gene families, suggesting a compensatory relationship between these two routes of evolution. This pattern is most pronounced in evolutionarily dynamic functional categories of genes, in new genes families compared to ancestral ones and in individual genomes compared to groups of related genomes analyzed as a whole. Putting these observations together, we propose a model of genome evolution (**Figure 3)**. Evolution of a large fraction of proteins involves neither LCR nor paralogy, forming the major attractor at the origin of the LCR-PAR plot. However, in many other cases, greater evolutionary flexibility appears to be essential whereby new functions require new raw material to evolve. Such dynamic genes escape the attractor at the origin via two main routes of evolution, the increase in LCR that is characteristic of fast gene evolution on a short time scale, and increase in gene paralogy that is typical of functions evolving on a longer time scale. As both mechanisms also impose genetic burden on the organism, we conjecture the existence of another (and wider) attractor characterized by optimal values of LCR and PAR. The dynamic interplay between these two routes of evolution eventually results in the observed inverse relationship between LCR and gene paralogy.

**Figure 3.**
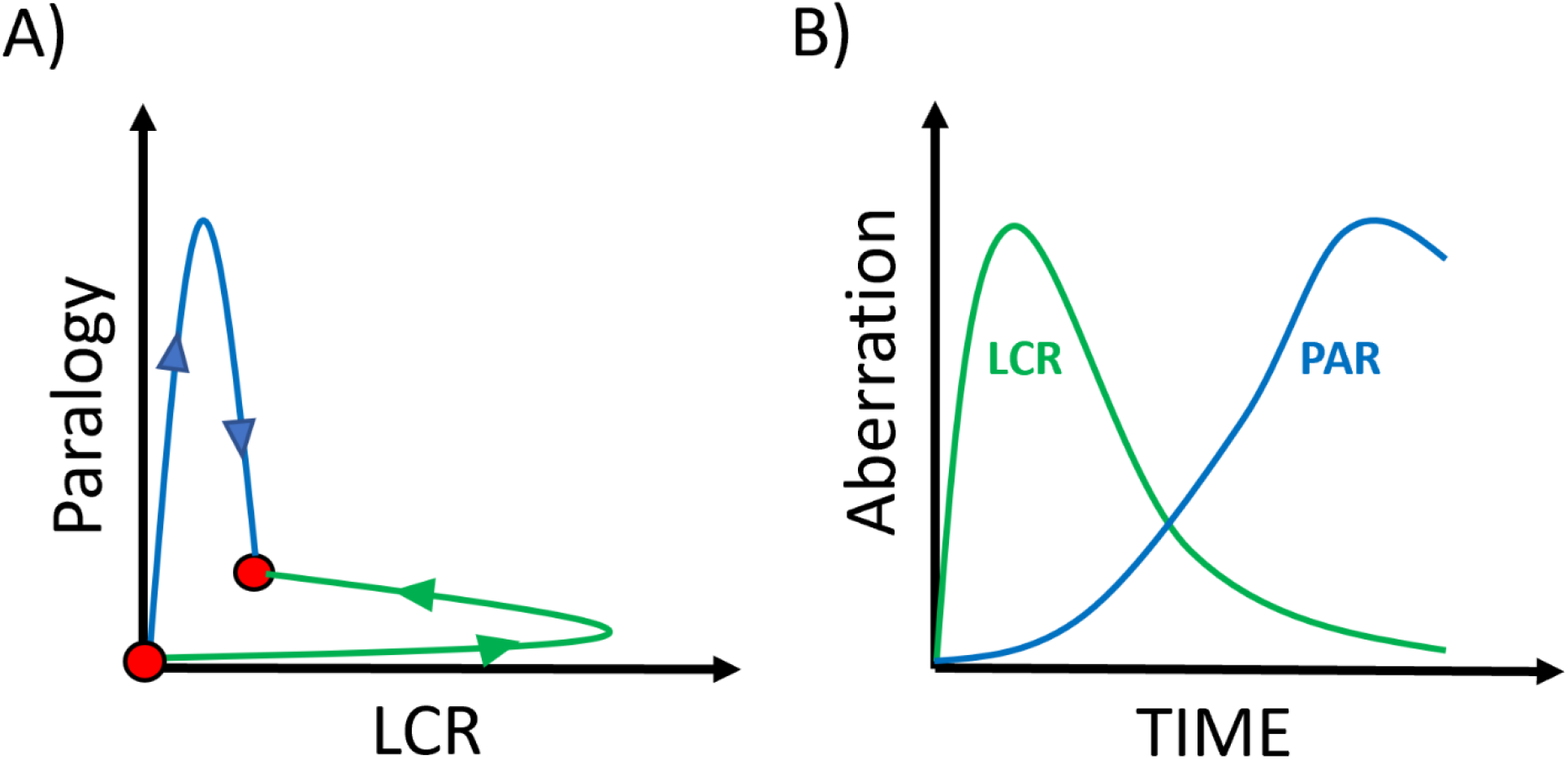
Model of the compensatory dynamics of low complexity regions and gene paralogy in species evolution. (**A**) There is an attractor of functions evolving without the need for LCR or PAR (i.e., LCR=0, PAR=1, red origin), which represents the null solution of evolution by point or small-scale mutations in the respective protein-coding genes. However, this mode is insufficient for the evolution of many biological functions, which resort to additional genomic mechanisms. One solution, dominant in ancestral and slowly evolving functions, is to gain new gene copies (PAR>1; blue curve), whereas the other, dominant in recent and presumably fast evolving functions, is to expand low complexity regions (LCR>1; green curve). The existence of these two distinct routes manifests as an inverse relationship between LCR and PAR across different COGs and suggests the existence of a second attractor with optimal levels of LCR and PAR (second red circle), leading to the observed scatter of points (cf. Fig 1A). (**B**) An illustration of the fast transient dynamics of LCR (green), rising and dominating short-term protein evolution, but subsequently eliminated by purifying selection, and supplanted by gene paralogy (blue).

The dynamics of LCRs observed here is consistent with the previous observations on the roles of such regions in fast evolution of new functions, such that they are more abundant in relatively young proteins^15,16^. With time, these regions accumulate mutations and diverge, so that the overall LCR reduces. The reduction in LCR over time by deletion is further driven by purifying selection, once functions become fixed and conserved, explaining the overall low LCR in ancient proteins, in contrast to young proteins that are subject to more variable selection pressures^49^.

Compensatory relationships between short-term and longer-term mechanisms are likely to represent a universal feature of the evolutionary process in all kinds of biological contexts. Analysis of the dynamics of different types of genome aberrations in cancer revealed patterns similar to that observed here for the evolution of prokaryotes^39,40^. Short repeats instability is prominent at early stages of tumor evolution, but subsequently declines and is supplanted by point mutations, and then, by larger scale aberrations, such as copy number variation and chromosomal aneuploidy. In experimental evolution of poxviruses, a similar compensation was observed between rapid gene amplification providing initial adaptation followed by fixation of point mutations leading to the same effect and allowing elimination of amplified genes, apparently, under the selective pressure to reduce the burden incurred by extra gene copies on genome replication^50^. It appears likely that many more levels of compensatory interplay between evolutionary processes remain to be discovered, in accordance with the general concept of recapitulation of evolutionary mechanisms at different temporal scales and different levels of biological organization^51^.

## Methods

### Datasets

we analyzed a dataset of 8 groups of species, 4 archaeal: Sulfolobales (n=52), Thermococcales (n=42), Methanosarcina (n=41), Haloferacales (n=37), and 4 bacterial: Deinococcales (n=33), Flavobacteriales (n=50), Rhodococcus (n=53), Paenibacillaceae (n=66) that we have curated and published recently^52^. Briefly, for each group of species, the set of local clusters of orthologous genes (COGs) was delineated, which defines for each COG a set of distinct proteins associated with it, across the genomes in the respective group of species. Using 16S rRNA we build species tree and assigned for each COG a time signature: Ancestral (T=0) if the COG exists in all the genomes comprising a group of species, or Recent/Young (T=1) if the COG is represented in only one leaf. For a complete description of the datasets reconstruction, see^52^.

### Measures of low complexity and paralogy

the level of gene paralogy (PAR) was defined by the number of distinct genes (associated with a COG) across the genomes in a group of species, divided by the number of genomes in which the COG is represented. Thus, PAR is always ≥ 1. The extent/coverage of Low Complexity Regions (LCR) was estimated using a dedicated algorithm to extract repeats, which identifies all short *K*-mers that recur beyond random and process their recurrences and locations, as described below. The method was applied to the nucleotide sequences of the datasets.

### LCR detection pipeline (LCRFinder)

In random sequences, the probability of identifying a motif of length *k* that recurs more than *n* time in a sequence of length *L* is determined by the binomial distribution, 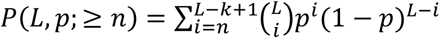,, where *p* is the probability of selecting a *k*-mer over an alphabet *A*, such that *p* = 1/*A*^*k*^. We define non-random recurrences of a *k*-mer as those that recur with *P* <10^−6^, locally (that is within a window of 1000 characters), using a default parameterization of K=1-6 for nucleotide sequences and K=1-3 for amino-acid sequences. Thus, for example, for nucleotide sequences (*A* = 4), and using *k* = 6 (hexamers), this definition translates into the search of non-random recurrence of hexamers that occur at least 6 times within 1000 bp, at least 5 times within 500 bp, at least 4 times within 200 bp and at least 3 times within 40 bp. To extract LCR, we identify all non-random recurrences with interval distance (*I*) between consecutive recurrences of the same motif smaller than the motif length *k* (i.e., *I* ≤ *k*) – that is all overlapping instances of the same motif, since they represent pure tracks of words. For example, using *k* = 6 in the sequence AAAAAAAAA, the hexamer AAAAAA recurs 4 times, with *I* = 1 between consecutive recurrences, capturing runs of nucleotides. Similarly, in the sequence ATATATATATA the dinucleotide tandem repeats are captured by the hexamers ATATAT and TATATA, each recurring 3 times, with distance interval *I* = 2 (recurring twice each), and so forth up to *I* = 6. These instances represent pure long tracks. To ensure that all non-random patterns are identified, this procedure is done for all *k* up to 6 (i.e., *k* = 1. .6). LCRs that are separated from each other by less than *k*, are merged into a single region. Using the default parametrization, LCRs in nucleotide sequences include the conventionally defined regions of microsatellites (i.e., tracks of units composed of a few bp, typically 1-5 bp).

Because integration over the probability *P* is implemented in practice, to best cover the repertoire of LCRs, the algorithm does not require any choice of parameters (such as window size, signal detection thresholds or similarity scores), except for *k*, which separates LCRs from longer and more complex repetitive sequences (if the repeat unit is not LCR in itself) which form different classes of repeats than LCRs. The algorithm allows as well to obtain different resolutions of LCRs (from pure tracks to diverged sequences). Examples of the application of the LCR pipeline to protein sequences are shown in ***SI*** (**Figs. S8-S10**). The algorithm is developed in MATLAB and is available through Github (https://github.com/erezpersi/LCRFinder).

## Supporting information

List of COGs in 8 groups of species

