## Supplementary Information for "Compensatory Relationship between Low Complexity Regions and Gene Paralogy in the Evolution of Prokaryotes"

#### This File includes

Figs. S1-S10

#### Other supplementary materials for this manuscript include the following:

**Dataset S1** – List of Cogs for the 8 groups of species. Provided as **XLSX Table**.

**Software Package S1** - The computational Pipeline, implemented in MATLAB, to extract Low complexity Regions (LCR) from nucleotide and amino acids sequences. The pipeline can be downloaded via **GitHub** (https://github.com/erezpersi/LCRFinder). Follow the READ_ME file and/or type 'help run_LCRFinder' in Matlab command line to execute examples of processing amino-acid and/or nucleotide sequences. Executing run_LCRFinder.m also prints the output (i.e., the detected LCRs) to a file (numbered), in the format (loc_start) seq (loc_end), for each input sequence.

### Figures

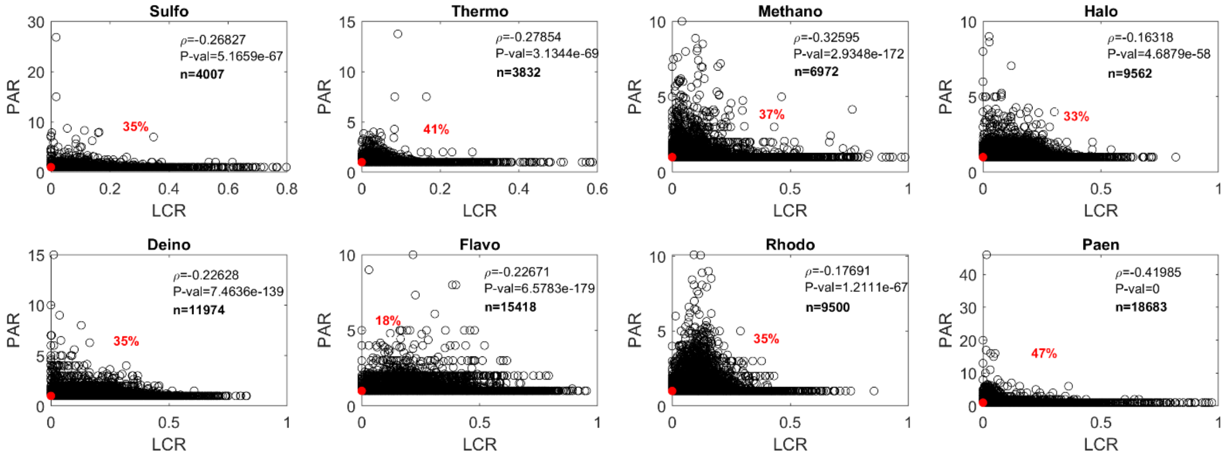


**Figure S1.** The scatter plot and Spearman correlation between LCR and PAR of local COGs on the set of 8 groups of species: 4 archaea 4 (upper panel), and 4 bacteria (lower panel). Same a Figure 1A, for each of the 8 groups of species. Percentage in red indicate the no. of cogs at the origin (i.e., LCR=0 and PAR=1), and the actual no. of cogs excluding the origin are shown, used for the correlation analysis. **The Inverse shape of the scattered point is evident across all groups of species.**

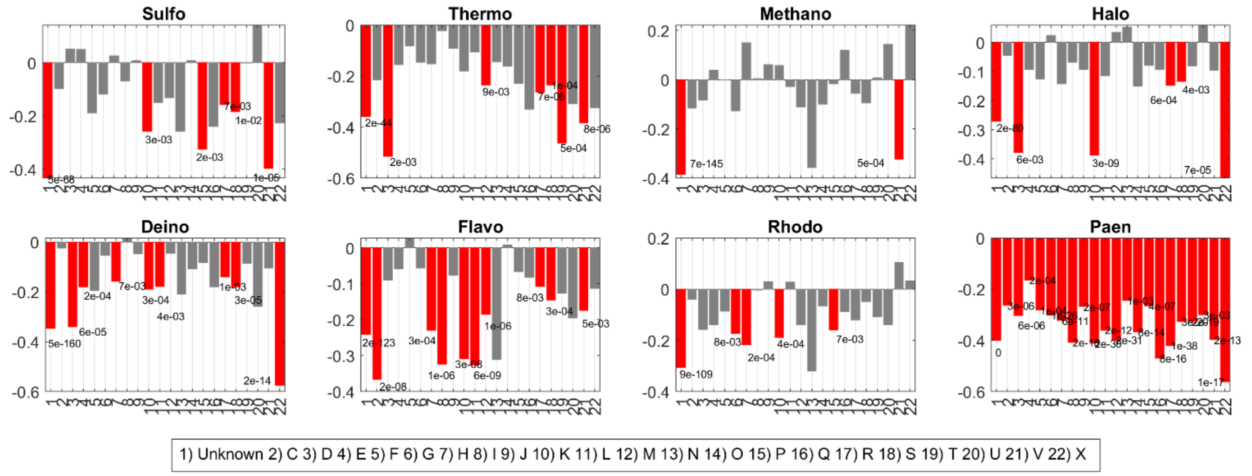


**Figure S2.** The Spearman correlation between LCR and PAR of COGs belonging to different functional categories. Bars marked in red have P-values < 0.01. Same a Figure 1B, for each of the 8 groups of species: 4 archaea 4 (upper panel), and 4 bacteria (lower panel). **The negative correlation between LCR and PAR is apparent across most of the biological functions, and across all groups of species.**

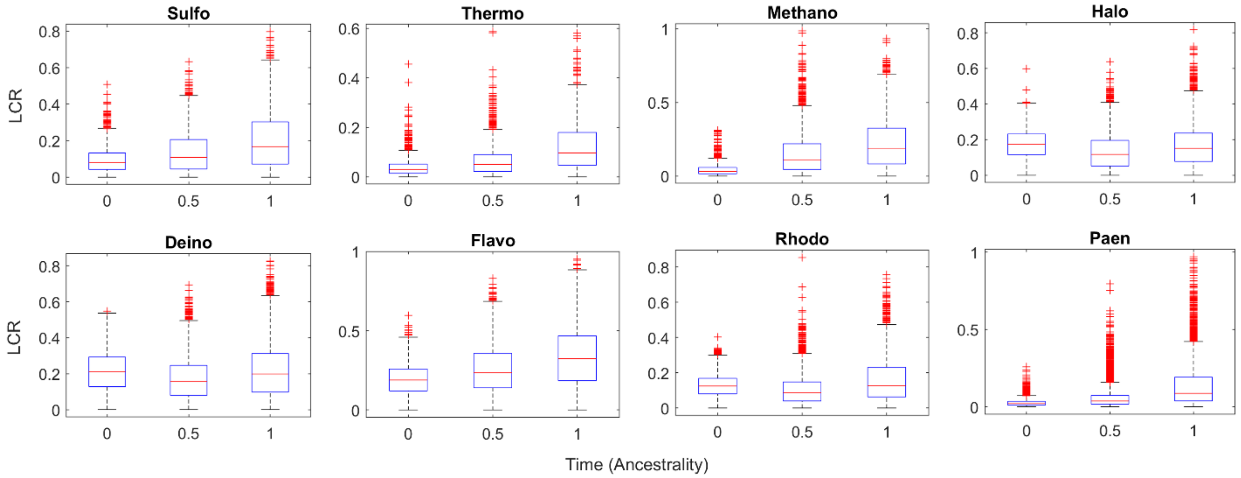


**Figure S3.** LCR level across time (age) of COGs (Ancestral, T=0, Intermediate, T=0.5, Recent, T=1, based on the species tree of each group of species) for the set of 8 groups of species: 4 archaea 4 (upper panel), and 4 bacteria (lower panel). Same a Figure 1C, for each of the 8 groups of species. **There is clear evidence for larger LCR at T>0 that it is at T=0 across most species.** Note that even when levels are comparable (e.g., Halo, Deino, Rhodo), the tails of the distributions are heavier at T>0 than the tail of distribution at T=0.

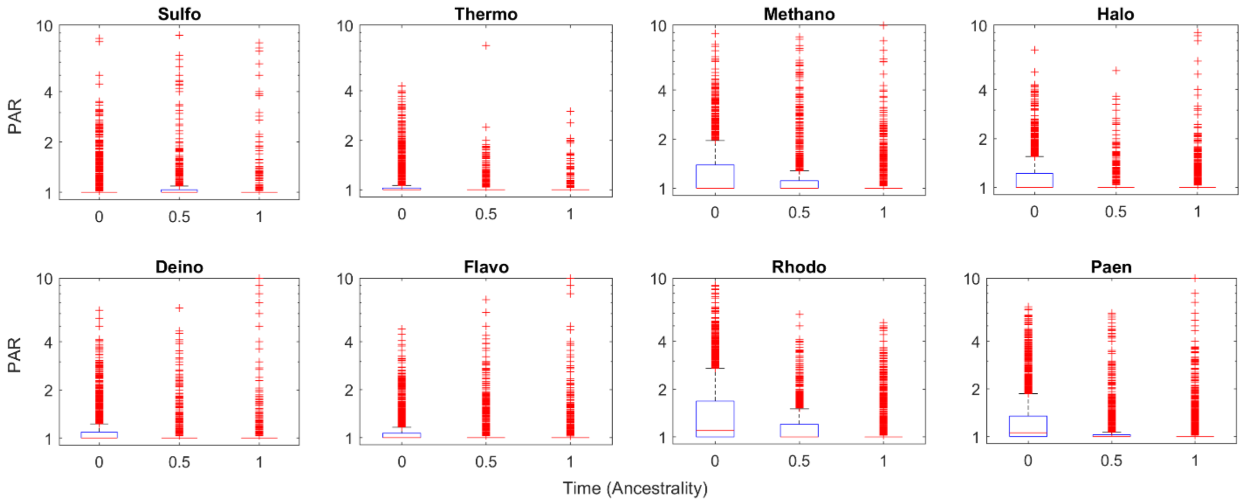


**Figure S4.** Level of Paralogy (PAR) across time (age) of COGs (Ancestral, T=0, Intermediate, T=0.5, Recent, T=1) for the set of 8 groups of species: 4 archaea 4 (upper panel), and 4 bacteria (lower panel). **There is apparent tendency for larger PAR values in locally ancestral COGs (T=0) than the values in younger COGs (T>0).** For clarity, the y-axes are shown in log scale and values of PAR > 10 are not displayed.

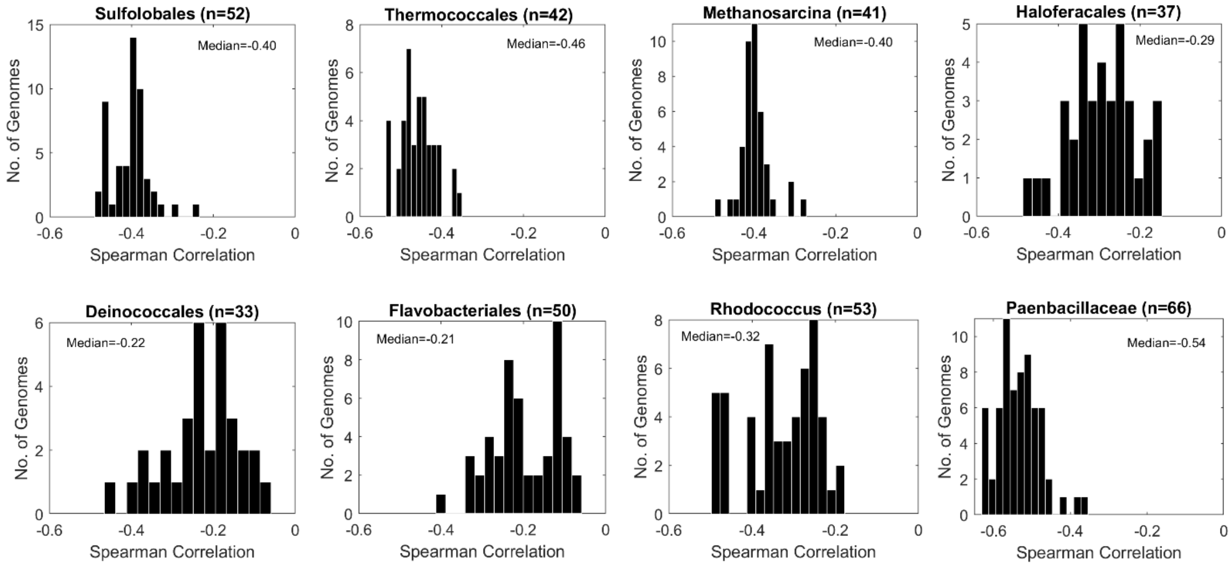


**Figure S5.** The distribution of the Spearman correlation between LCR and PAR, when estimated at the single genome level. The distribution across all the genomes in each of the 8 group of species is shown (4 archaea, upper panel; 4 bacteria, lower panel). The medians of the distribution are provided.

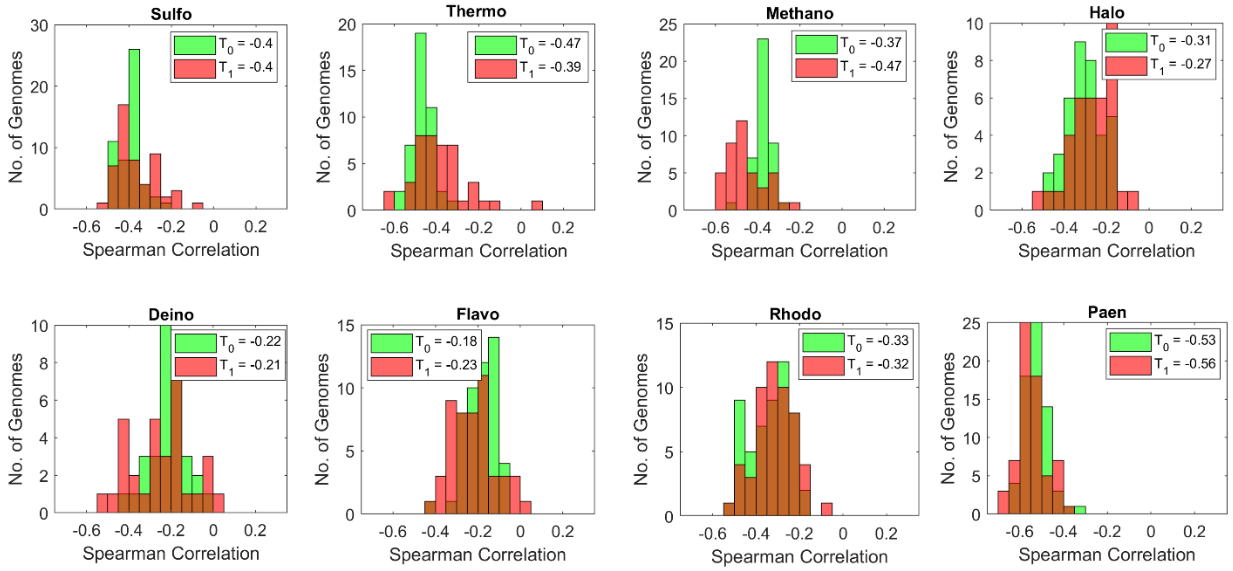


**Figure S6.** The lack of break in correlation between locally ancestral COGs (T=0) and recent/young COGs (T=1), as derived from the local species tree in each of the 8 groups of species. This contrasts with the case when LCR and PAR are estimated based on the gene x genomes full matrix (cf. **Figure 1D**). The reason for this turns out to be that LCR levels (and not PAR levels) are much high in single genomes (see **Figure S7**)

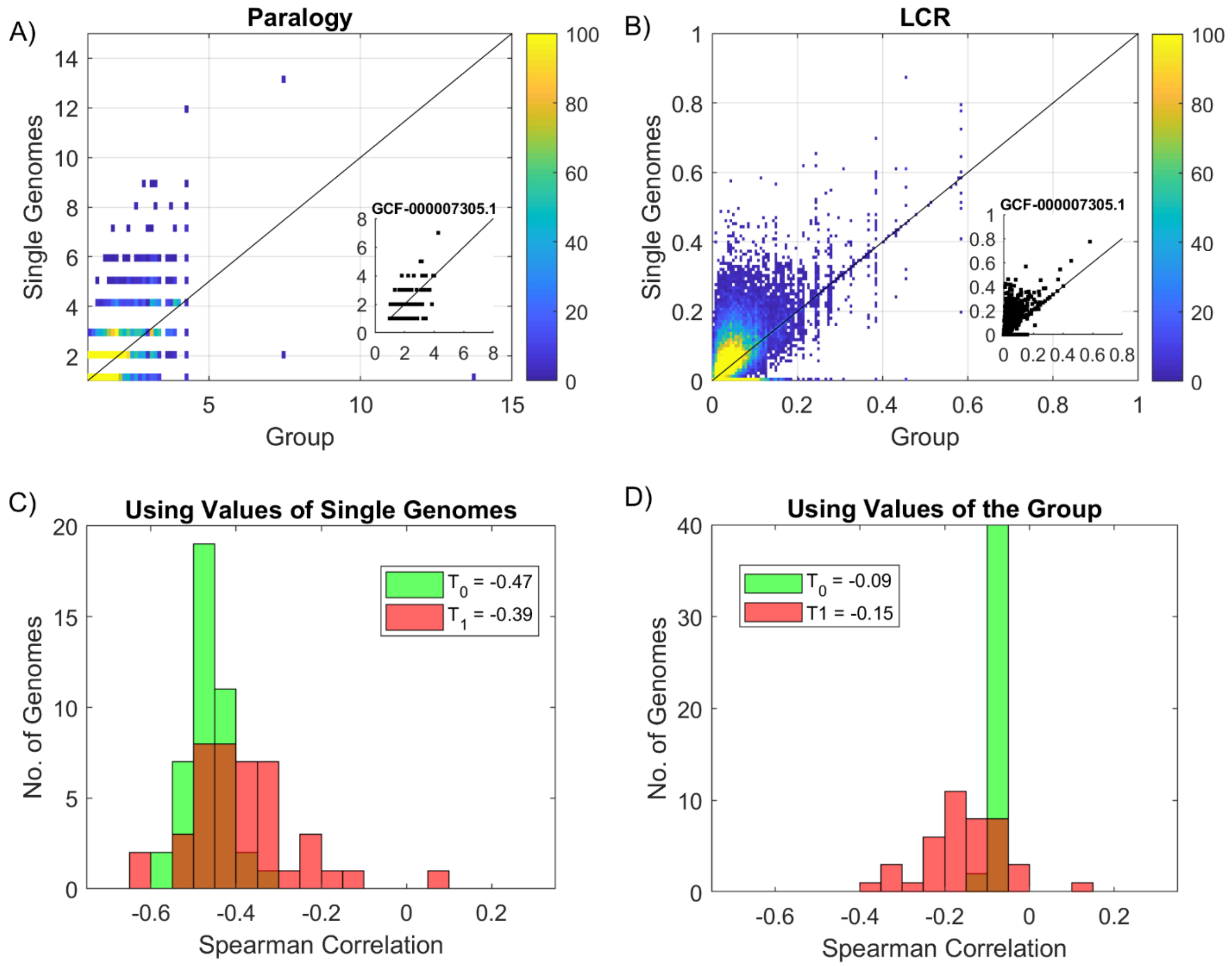


**Figure S7. The origin for strong negative correlation between LCR and PAR in single genomes** is not the extent of Paralogy (**A**) but rather the much larger extent of LCR (**B**) in single genomes superimposed (y-axis) compared with when these measures are calculated for all proteins associated with these COGs across all genomes in the respective group of species (x-axis). Thermococcales group of species (n=42) are used in this figure. Lines denotes linear relationship. Color denote the density of COGs. Insets show the case of particular selected genome (GCF_000007305.1 - Pyrococcus furiosus DSM 3638) (**C**) and (**D**) shows respectively the strong correlation across single genomes, in both time scales, Ancestral ($T_0$) and Recent ($T_1$) when LCR and PAR are calculated at the single gnome level (C), and the (relative) break of this correlation (like the effect shown in Figure 1 of the main text) when they are calculated from the full matrix of genes x genomes in a group of species (D).

MESSAKMESGGAG**QQPQPQPQQ**PFLPPAACFFAT**AAAAAAAAAAAAA**QSA**QQQQQQQQQQQQ**APQLRPAADGQPSGGGH
KSAPKQVKRQRSSSPELMRCKRRLNFSGFGYSLP**QQQPAA**VARRNERERNRVKLVNLGFATLREHVPNGAANKKMSKVE
TLRSAVEYIRALQQLLDEHDAVSAAFQAGVLSPTISPNYSNDLNSMAGSPV**SSYSS**DEGSYDPLSPEEQELLDFTNWF

**Figure S8. LCR pipeline applied to amino-acid sequences**. Example of LCR coverage of the protein **ASCL1** (Swissprot ID: **P50553**): a transcription factor that plays a key role in neuronal differentiation, acts as a pioneer transcription factor, accessing closed chromatin to allow other factors to bind and activate neural pathways. In Grey is the LCR coverage reported by Swissprot as compositional bias of Ploy-A (33-47) and Ploy-Q (51-62). In color is the LCR coverage by the method introduced in this study using K=1-3 (the default parametrization for amino-acid sequences): green represents pure tracks at K=3, and blue denotes the additional coverage identified by accounting for irregular recurrences at K=1 and K=2.

MSDASLRSTSTMERLVARGTFPVLVRTSACRSLFGPVDHEELSRELQARLAELNAEDQNRWDYDFQQDMPLRGPGRLQWTEVD
SDSVPAFYRETVQVGRCRLLLAPRP**VAVAVAVSPP**LEPAAESLDGLEEAPEQLPSVPV**PAPASTPPPVPVLAPAPAPAPAPVA**
**APVAAPVAVAVLAPAPAPAPAPAPAPAPVAAPAPAPAPAPAPAPAPAPAPDAA**PQESAEQGANQGQRGQEPLADQLHSGISGR
P**AAGTAAA**SANGAAIKKLSGPLISDFFAKRKRSAPEKSSGDVPAPCPSPSAAPGVGSVEQTPRKRLR

**Figure S9. LCR pipeline applied to amino-acid sequences**. Another example of LCR coverage, here applied to the protein **CDKN1C** (Swissprot ID: **P49918**): Cyclin-dependent kinase inhibitor 1C. Potent tight-binding inhibitor of several G1 cyclin/CDK complexes (cyclin E-CDK2, cyclin D2-CDK4, and cyclin A-CDK2). In Grey is the coverage reported by Swissprot as 9 repeats of the motif PAPA. In color is the LCR coverage by the method introduced in this study using K=1-3 (the default parametrization for amino-acid sequences): green represents pure tracks at K=3, and blue denotes the additional coverage identified by accounting for irregular recurrences at K=1 and K=2.

MKIITYFCIWAVAWAIPVPQSKPLERHVEKSMNLHLLARSNVSVQDELNASGTIKESGVLVHEGDRGRQENTQDGHKGEGNGSKWAEVGGKSFST
YSTLANEEGNIEGWNGDTGKAETYGHDGIHGKEENITANGIQGQVSIIDNAGATNRSNTNGNTDKNTQNGDVGDAGHNEDVAVVQEDGPQVAGSN
NSTDNEDEIIENSCRNEGNTSEITPQINSKRNGTKEAEVTPGTGEDAGLDNSDGSPSGNGADEDEDEGSGDDEDEEAGNGKDSSNNSKGQEGQDH
GKEDDHDSSIGQNSDSKEYYDPEGKEDPHNEVDGDKTSKSEENSAGIPEDNGSQRIEDTQKLNHRESKRVENRITKESETHAVGKSQDKGIEIKG
PSSGNRNITKEVGKGNEGKEDKGQHGMILGKGNVKTQGEVVNIEGPGQKSEPGNKVGHSNTGSDSNSDGYDSYDFDDKSMQGDDPNSSDESNGND
DANSESDNNSSSRGDASYNSDESKDNGNGSDSKGAEDDDSDSTSDTNNSDSNGNGNNGNDDNDKSDSGKGKSDSSDSDSSDSSNSSDSSDSSDSD
SSDSNSSSDSDSSDSDSSDSSDSDSSDSSNSSDSSDSSDSSDSSDSSDSSDSKSDSSKSESDSSDSDSKSDSSDSNSSDSSDNSDSSDSSNSSNS
SDSSDSSDSSDSSSSSDSSNSSDSSDSSDSSNSSESSDSSDSSDSDSSDSSDSSNSNSSDSDSSNSSDSSDSSNSSDSSDSSDSSNSSDSSDSSD
SSNSSDSSDSSDSSDSSDSSNSSDSNDSSNSSDSSDSSNSSDSSNSSDSSDSSDSSDSDSSNSSDSSNSSDSSDSSNSSDSSDSSDSSDGSDSDS
SNRSDSSNSSDSSDSSDSSNSSDSSDSSDSNESSNSSDSSDSSNSSDSDSSDSSNSSDSSDSSNSSDSSESSNSSDNSNSSDSSNSSDSSDSSDS
SNSSDSSNSSDSSNSSDSSDSNSSDSSDSSNSSDSSDSSDSSDSSDSSDSSNSSDSSDSSDSSDSSNSSDSSNSSDSSNSSDSSDSSDSSDSSDS
SDSSDSSDSSNSSDSSDSSDSSDSSDSSDSSDSSDSSESSDSSDSSNSSDSSDSSDSSDSSDSSDSSDSSDSSDSSNSSDSSDSSDSSDSSDSSN
SSDSSDSSESSDSSDSSDSSDSSDSSDSSDSSDSSDSSNSSDSSDSSDSSDSSDSSDSSDSSDSSDSSDSSDSSDSSDSSDSSDSSDSSDSNESS
DSSDSSDSSDSSNSSDSSDSSDSSDSTSDSNDESDSQSKSGNGNNNGSDSDSDSEGSDSNHSTSDD

**Figure S10. LCR pipeline applied to amino-acid sequences**. Another example of LCR coverage, here applied to the protein **DSPP** (Swissprot ID: **Q9NZW4**). encodes a member of the small integrin-binding ligand N-linked glycoprotein (SIBLING) family of proteins. The encoded preproprotein is secreted by odontoblasts and proteolytically processed to generate two principal proteins of the dentin extracellular matrix of the tooth, dentin sialoprotein and dentin phosphoprotein. In Grey is the coverage reported by Swissprot. In color is the LCR coverage by the method introduced in this study using K=1-3 (the default parametrization for amino-acid sequences): green represents pure tracks at K=3, and blue denotes the additional coverage identified by accounting for irregular recurrences at K=1 and K=2.
